# ABAG-Rank: Improving Model Selection of AlphaFold Antibody–Antigen Complexes by Learning to Rank

**DOI:** 10.64898/2026.03.17.712376

**Authors:** Matteo Tadiello, Marko Ludaic, Vsevolod Viliuga, Arne Elofsson

## Abstract

**Motivation:** AlphaFold has transformed structural biology with an unprecedented accuracy in modeling protein structures and their interactions with biomolecules, with AlphaFold3 (AF3) achieving state-of-the-art performance. However, AF3 and other methods often struggle to accurately predict the structure of protein complexes that lack strong co-evolutionary information, such as antibody-antigen (Ab-Ag) complexes. One of the fundamental issues is that AF3 often generates accurate predictions, but fails to reliably distinguish them from the much larger set of incorrect ones.

**Results:** To address this, we propose ABAG-Rank, a deep neural network that provides an efficient and robust solution for model selection of Ab-Ag interactions from a pool of structural ensembles predicted with AlphaFold. Built on the permutation-invariant DeepSets architecture, ABAG-Rank can process variable-sized ensembles of structural decoys and is directly applicable to prediction settings in which the number of candidates may vary. We train a model on a redundancy-reduced set of all known antibody-antigen complexes and find that simple geometric descriptors, along with confidence scores from AlphaFold, provide rich information about interface quality without requiring intensive physics-based calculations. Our experiments demonstrate that ABAG-Rank significantly outperforms AF3 internal scoring and the ranking performance of existing deep learning baselines.

**Implementation:** Source code can be found at: https://github.com/tadteo/ABAG-Rank

## Introduction

The release of AlphaFold2 (AF2) [1] has truly revolutionized the field of structural biology. By achieving unprecedented accuracy in predicting the three-dimensional structure of proteins from their one-dimensional amino acid sequences, AF2 has become a standard tool in almost all structural studies. Building on the foundations and best practices of AF2, AlphaFold3 (AF3) [2] was extended to model all-atom protein structures in complexes with metal ions, nucleic acids, small-molecule ligands, and other biomolecules. This computational breakthrough enabled researchers to infer protein functions and interactions prior to experimental validation, substantially reducing exploratory laboratory costs and efforts. However, accurately predicting molecular interfaces remains a grand challenge, particularly for complexes in which highly specific binding interactions determine recognition and affinity, and distinct co-evolutionary information may be too scarce or absent altogether.

Despite their overall high accuracy, the quality of the predictions produced with AF2/3 critically depends on the model’s internal confidence metrics, namely the predicted TM-score (pTM) and interface pTM (ipTM) scores [1]. These scores are used to rank candidate structures based on their predicted reliability. The internal ranking metrics, however, often fail to adequately differentiate between distinct predictions, often overestimating the quality of models with incorrectly predicted molecular interfaces, especially for specific subsets of proteins, such as antibody-antigen (Ab-Ag) complexes [3, 4]. Unlike general protein-protein interactions, Ab-Ag complexes stand out with their unique structural and functional complexity and the lack of co-evolutionary information [5]. Antibody binding is mediated by hypervariable loops (Complementarity-Determining Regions, CDRs), and particularly the H3 loop, which exhibits immense sequence diversity and conformational flexibility, obscuring strong structural prior needed for confident prediction. One of the most common issues is the placement of antibody chains at incorrect binding sites (epitopes) [3, 5]. AlphaFold often generates decoys in which the antibody is docked to the wrong epitope on the antigen - a region that may exhibit high geometric complementarity but in reality lacks biological relevance (Figure 1). These incorrect binding modes can nevertheless receive high pTM/ipTM confidence scores, distorting the ranking and effectively hallucinating a high-confidence interaction at an incorrect binding site. Consequently, binding poses that more closely align with the ground truth epitope can be overlooked if AF incorrectly scores them as lower-quality structures relative to these high-scoring false positives.

**Figure 1.**
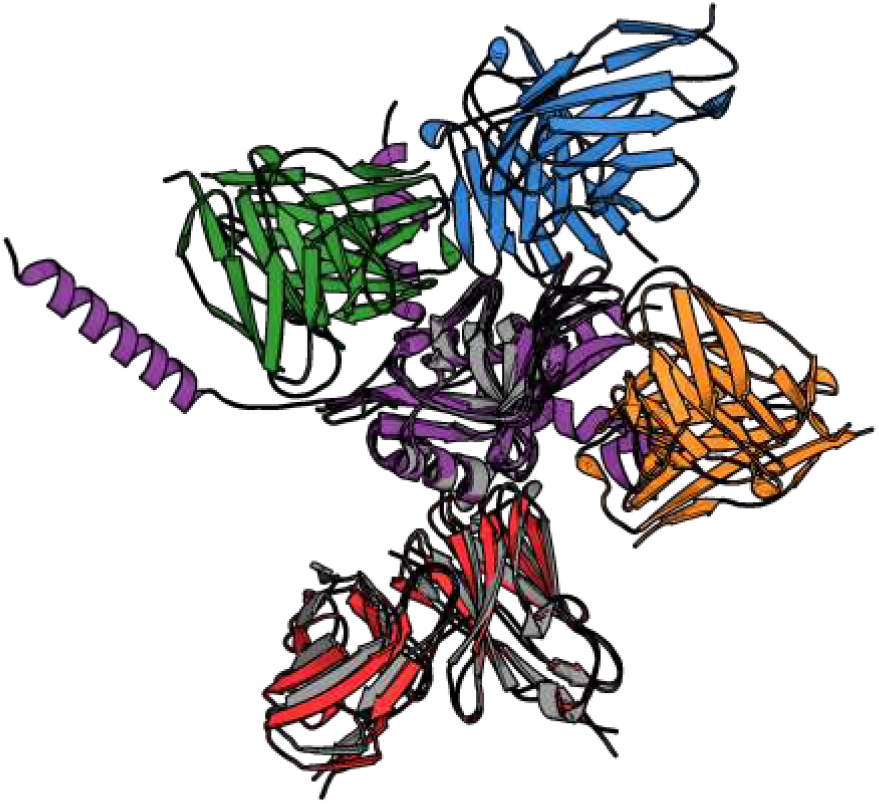
Structural diversity of AlphaFold3 predictions for an example antibody-antigen complex (PDB ID: 8URF). AlphaFold3 frequently places the antibody CDR loops at distinct, non-overlapping interfaces on the antigen. Although many of these binding modes might appear geometrically plausible, they are often biologically incorrect. The antigen
is shown in purple, the reference ground-truth structure in gray, and different predicted antibody poses in green, blue, orange, and red.

**Figure 2.**
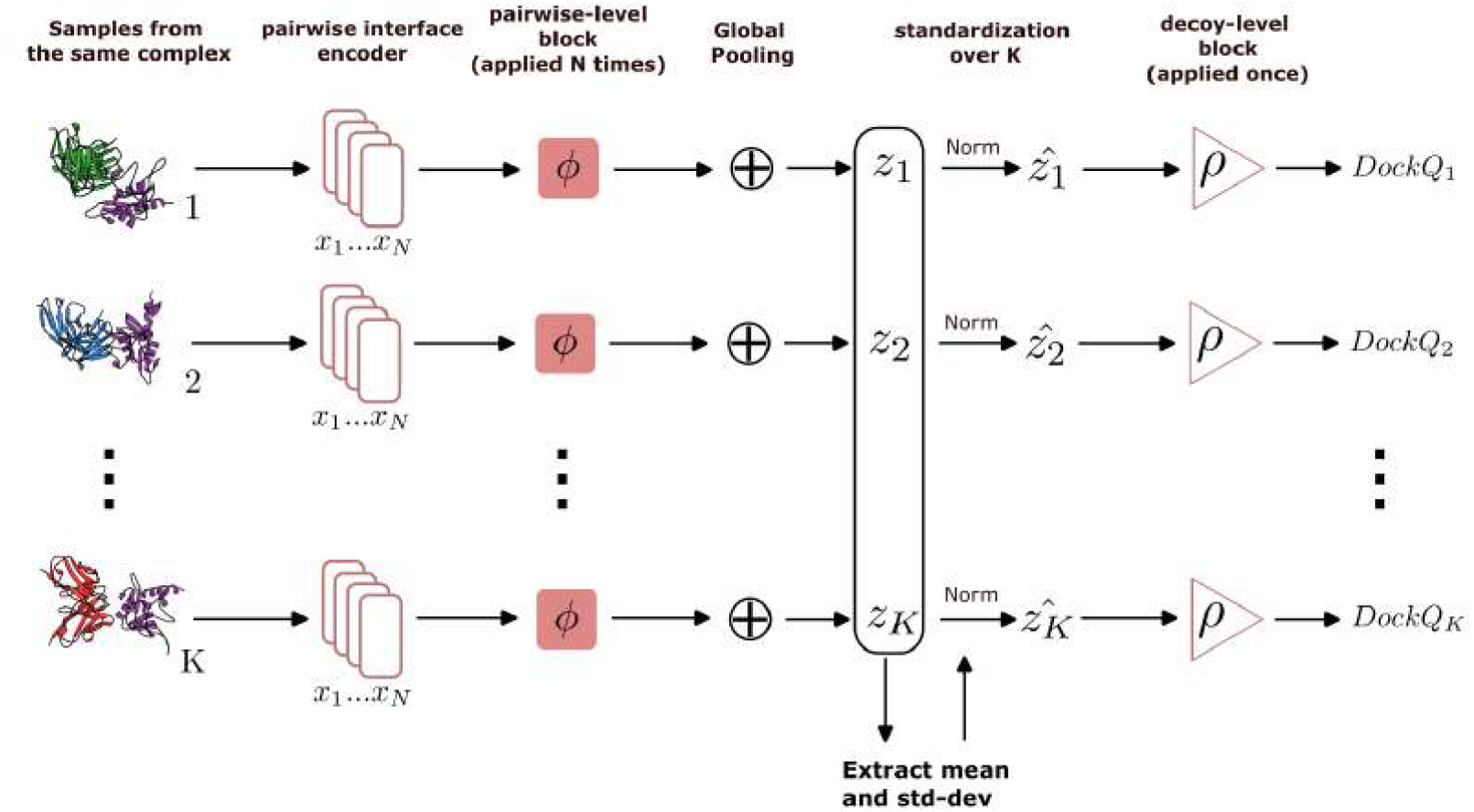
ABAG-Rank Architecture. The model processes a variable-sized set of *K* decoys per complex. (A) For each decoy, inter-chain residue pairs are encoded with geometric features (PAE, Distance, Residue type) and optionally evolutionary context (ESM embeddings) [18]. (B) These pair features are processed by a shared encoder *φ* and aggregated via concatenated statistics (Mean, Max, Attention, interface-length). (C) The resulting global descriptor is standardized over the K samples for complex b. (D) the normalized embedding is passed to a decoder ρ to predict the interface quality score.

One of the most established strategies for antibody-antigen structure prediction is extensive sampling; this approach trades computational cost for a higher probability of accurate predictions. AFsample [6], for instance, applies aggressive dropout-based stochastic inference with AF2 to generate large and diverse structural ensembles per protein complex, while MassiveFold [7] extends this strategy to large-scale computational clusters, significantly reducing computation time from days to hours. However, the persistence of unreliable confidence-based ranking renders ranking fidelity a fundamental limitation, such that even with aggressive sampling strategies, structurally more accurate solutions may still be overlooked if they are assigned lower confidence scores.

Aiming to improve on the heuristic confidence estimates of AF2/3, several alternative interface quality metrics have been been proposed. For example, pDockQ [8] and pDockQ2 [9] directly use PAE and pLDDT outputs of AF2 and map them to DockQ-like [10] scores through a simple sigmoid fit, resulting in improved interface quality estimation for protein complexes. Other approaches, such as actifpTM [11] and ipSAE [12], modify (i)pTM and PAE more explicitly: actifpTM reweights residue pairs using predicted distance probabilities such that interacting residues contribute more strongly than non-contacting ones, thereby reducing length-dependent bias from non-interacting regions. ipSAE, in turn, restricts the calculation to inter-chain residue pairs with reliably low PAE and rescales the score accordingly to reduce the inflated confidence. Despite the refinements, such purely AF confidence-based alternatives conceptually remain limited in their generalizability, particularly for antibody-antigen complexes, where AlphaFold confidence metrics alone are often insufficient to distinguish correct from incorrect binding modes. Alternatively, several deep learning-based interface scoring approaches, such as EuDockScore [13] and GdockScore [14] have been proposed. However, their applicability to antibody-antigen complexes remains unclear, as they have not been systematically benchmarked in this setting. The only scoring model specifically developed for Ab-Ag interactions is the recently released DeepRank-Ab [15], an equivariant graph neural network that predicts DockQ from a large set of biophysics-based interface descriptors. Because these features must be computed with HADDOCK [16], the method introduces external dependencies and is comparatively inefficient.

This work proposes **ABAG-Rank** - a deep neural network trained to rank the quality of predicted Ab-Ag models and designed to address the limitations of both heuristic AlphaFold-derived confidence scores and existing deep learning-based interface scoring approaches. Leveraging the DeepSet architecture [17], the model treats each decoy ensemble as a permutation-invariant, variable-sized set rather than as a collection of isolated instances, which is well suited to the practical setting where the number of predictions per complex varies and the structural diversity within an ensemble can range from highly heterogeneous to nearly collapsed around a single structural solution. By integrating regression and ranking objectives into a composite loss, ABAG-Rank is trained to resolve subtle quality differences between alternative binding modes within the same Ab-Ag complex, even in the presence of highly skewed DockQ distributions and strong complex-to-complex variability. In addition, we introduce a redundancy-reduced dataset generation strategy and show that a smaller but statistically diverse training set is sufficient to learn a robust ranking model. Our experiments show that ABAG-Rank consistently outperforms both AF3 and DeepRank-Ab in ranking interface quality and in retrieving the highest-quality models from predicted structural ensembles. We further demonstrate that AF3-derived confidence scores, particularly iPTM and inter-chain Predicted Aligned Error (PAE), together with pairwise interface distances and ESM2 protein language model embeddings [18], provide an informative basis for accurate interface quality assessment.

## Method

In the following section, we describe the dataset construction strategy, model architecture, and core design choices underlying ABAG-Rank. In order to train a model to rank the quality of the Ab-Ag complexes, we first generate a diverse dataset of Ab-Ag models by combining AF3 predictions with careful curation. Second, we develop a dedicated ranking component that complements the native AF3 confidence metrics. AF3 can produce multiple models for a given complex by varying the random seed or the number of diffusion timesteps. Our goal is to provide an interface quality predictor to rank models reliably, especially when multiple predictions of the same complex are available, and improve the model’s ability to distinguish good from bad interfaces.

### Dataset Preparation

Using the Structural Antibody Database (SAbDab) [19] as our initial source of antibody-antigen complexes, we constructed an Ab-Ag dataset designed to maximize diversity in interface quality and Predicted Aligned Error (PAE) matrices as well as structural diversity. We used data points (structure decoys) for training and testing, using a cutoff date approach as used in AlphaFold. Using the following split:

- **Training:** all SAbDab entries till 2024/09/30
- **Test:** SAbDab entries from 2024/09/30 to 2025/06/01

For each complex, we run inference with AF3 using 10 random seeds and generate in total 50 structural decoys (5 samples per seed). For each of the decoys, we retrieve the full *N* × *N* Predicted Aligned Error (PAE) matrix (*N* : number of residues in a complex), along with the model’s internal confidence scores: pTM and ipTM, and the composite ranking score. To capture explicit geometric constraints, we compute pairwise C*α* distance matrices and derive per-residue features from them (i.e. each residue’s distance to all other residues). Since our focus is to quantify the interface quality, special attention is paid to inter-chain interactions; indices corresponding to antibody-antigen interface residues are extracted to emphasize the model’s focus on the binding interface.

To generate ground truth labels for training, we compare each generated decoy against its experimentally determined reference structure in the SabDab database. Given that antigens may comprise multiple polypeptide chains, we implement a logical chain-merging step prior to scoring. All antigen chains are merged into a single “Ag” entity, and antibody heavy/light chains are merged into an “Ab” entity. The structural quality is then assessed using DockQ [10].

Decoys where metric computation fails (e.g., due to alignment timeouts or missing chains) are filtered out to maintain dataset integrity.

Prior to redundancy reduction, we compute ensemble-level statistics, namely the column-wise mean, median, and standard deviation of the inter-chain PAE and *Cα* distances, across all generated decoys. These global context features are calculated on the full, unfiltered ensemble to ensure they accurately reflect the aleatoric uncertainty of AF3 predictions. Preservation of these statistics allows the model to assess an individual decoy’s confidence relative to the true population distribution, even after redundant samples are pruned.

We define redundancy based on three criteria: structural quality similarity (DockQ and RMSD) and confidence profile similarity (PAE distribution). First, we establish an adaptive PAE similarity threshold, *τ*_PAE_, for each complex. This is derived by randomly sampling 500 decoy pairs, computing the Wasserstein distance between their flattened PAE vectors, and setting *τ*_PAE_ to the 25th percentile of these distances. The DockQ difference threshold is fixed at *τ*_DockQ_ = 0.01 the RMSD difference threshold is fixed at *τ*_RMSD_ = 4 Å.

We then apply a greedy pruning algorithm to cluster similar decoys (Algorithm 2 is provided in Appendix S1). A sample is considered redundant (Algorithm 1) and removed only if it is sufficiently similar to an already selected sample in terms of PAE distribution (Wasserstein distance <*τ*_PAE_) *and* has a nearly identical DockQ score (difference <*τ*_DockQ_) and the RMSD of the sample is <*τ*_RMSD_. This ensures that we retain multiple conformations if they differ significantly in either geometry or model confidence.

The final dataset contains 4610 distinct complexes in the training and validation sets, with an average of 20 distinct samples per complex, and 1091 complexes in the test set, with an average of 19 distinct samples per complex.

### Model architecture and nested complex batching

We adopt a DeepSet-inspired architecture designed to process variable-sized sets of inter-chain residue pairs while preserving permutation invariance. The permutation-invariant architecture is essential to treat the interface features of Ab-Ag complexes as a set of individual pairs, focusing on the distance and properties and preventing the model from picking up unwanted biases stemming from residue ordering. Moreover, this architecture dynamically handles *N* pairs. By using global pooling, we collapse any number of residue-level features into a fixed-length “complex-level” embedding. This allows us to score any complex independent of the size. Moreover, PAE is inherently asymmetric; for this reason, (*i, j*) and (*j, i*) pairs have different values and both are processed by the model. In contrast to standard pooling, we employ an aggregation strategy that combines multiple permutation-invariant statistics with a global attention pooling head, allowing the model to capture both the average interface quality and sparse structural outliers within an interface. We restrict the set of pairs to those within a spatial cutoff of 12 Å (based on inter-chain *Cα* distance), treating pairs beyond this cutoff as non-interacting and thus excluded from the interface representation.

We propose a nested per complex batching. For each complex in the batch *B*, a nested tensor is computed by sampling *K* decoys for each complex *b* independently. Each training sample has hence the shape *X* ∈ ℝ ^*B*×*K*×*N*×*F*^ to reflect the underlying *set-of-decoys per complex* structure unique of the ranking problem, where *B* is the batch size, *K* is the number of decoys sampled per complex, *N* is the number of inter-chain residue pairs and *F* the number of features per residue pair. This helps to differentiate decoys from the same complex population. This hierarchical batching strategy encourages the network to learn *within-complex* relative ordering: the model observes diverse conformations in the same antibody–antigen context. This step is essential for the model to enforce the capability to differentiate interfaces for predictions of the same complex *b* using the information shred by each decoy *k*.

### Model input features

We hypothesize that much of the information required for accurate interface ranking is already implicitly encoded in (i) geometric features of the predicted interface and (ii) residue-level evolutionary signals captured by protein language model embeddings. Inter-chain *C*_*α*_ distances provide a direct description of geometric complementarity, while residue type encodings and ESM2 embeddings supply some learned biochemical context that can help distinguish biologically meaningful interactions from geometrically plausible artifacts. We further expect AlphaFold3 internal confidence scores to provide complementary information about the reliability of the predicted interface. Given the permutation-invariant architecture, positional encodings are included to restore sequential pair-specific information that would otherwise be lost in set-based processing.

Hence, for each residue pair (*i, j*) we construct a feature vector *x*_*ij*_ ∈ ℝ^*F*^ comprising:

- inter-chain *Cα* Euclidean distance;
- inter-chain predicted aligned error (PAE), centred by the complex-level mean;
- complex-specific mean interface PAE (global context);
- positional encoding based on a Cantor pairing of indices (*i, j*), normalized to [0, 1];
- residue type encoding (e.g., one-hot identity and/or ESM2-derived features).

Each pair vector is then processed independently by a shared encoder *φ* implemented as a multi-layer perceptron (MLP):

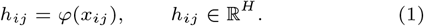

Given the set of encoded pair features 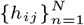 for a decoy, we use a statistical concatenation strategy with a global attention pooling head to aggregate the information. First, we compute standard statistics:

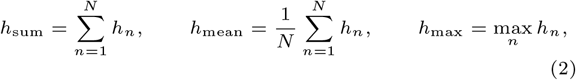

(where max is taken element-wise). To additionally capture sparse, high-importance residue pairs, we compute an attention-pooled summary.

We then form the globally pooled representation by concatenating attention pooling with the standard statistics, and append an explicit set-size feature:

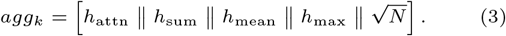

Where *N* is the number of interface residue pairs of a single decoy.

We then enforce the model to **explicitly represent intra-complex differences** using the information from the **nested complex batch**. For each element *b*, the aggregator normalizes the output using the mean and standard deviation across all *K* decoys for that specific complex in the nested batch.

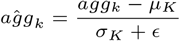

Where *µ*_*K*_ and *σ*_*K*_ are the mean and standard deviation of *agg* across the *K* decoys for complex

This strategy removes identity bias for the specific complex, i.e., predictions within the same complex can share similar interfaces. By normalising over the *K* decoys, you remove the “Complex Identity” and force the model to focus on the specific improvements of the decoy in a batch relative to the other decoys of the same complex. Each decoy’s final score is mathematically “aware” of its *K* peers. This enables the model to learn ranking even before it reaches the ranking loss function.

Finally, **aĝg**_**k**_ is normalized and passed to a decoder network *ρ* to predict a scalar score in logit space:

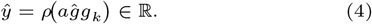

### Composite Loss Function

To optimize both (i) accurate quality prediction and (ii) consistent ranking of decoys *within each complex*, we minimize a composite objective:

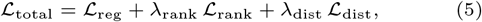

where *λ*_rank_ and *λ*_dist_ are fixed hyperparameters.

Regression loss (*ℒ*_reg_)

Let *y*_*b,k*_ ∈ ℝ denote the ground truth DockQ between the antibody and the antigen target in logit space for decoy *k* of complex *b*, and let *ŷ*_*b,k*_ be the model output. We use a Smooth-*𝓁*_1_ (Huber) loss in logit space:

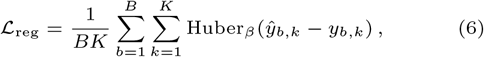

with robustness parameter *β*.

Soft Spearman Ranking (*ℒ*_rank_)

To directly optimize the ordering of decoys within each complex, we use a differentiable Spearman’s rank correlation loss. We first map logit outputs to DockQ space:

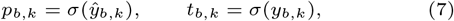

where *σ*(*·*) denotes the sigmoid.

For a score vector *s* ∈ ℝ^*K*^, we approximate the rank of element *k* via pairwise sigmoid comparisons with temperature *τ* :

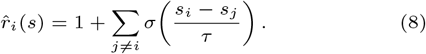

Let 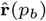 and 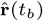 denote the soft-rank vectors for predictions and targets of complex *b*. We then compute a Spearman-style correlation by penalising squared rank discrepancies:

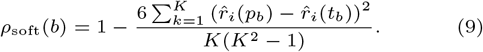

The ranking loss is defined as

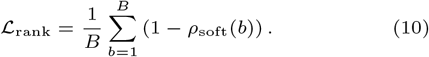

Since ranking is only meaningful when the *K* decoys for a complex exhibit non-trivial variation in quality (i.e. they do not fall strictly in the same structural cluster), we modulate the per-complex ranking loss by a continuous relevance weight based on the standard deviation of target scores:

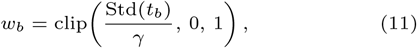

and use the gated loss

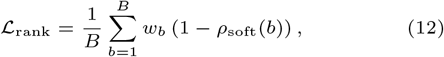

where *γ* is a scaling constant.

#### Distance Preservation (ℒ_dist_)

While *ℒ*_rank_ enforces correct ordering, it does not constrain the *magnitude* of predicted quality gaps. We therefore include a distance preservation objective operating on DockQ probabilities. For each complex *b*, we match pairwise absolute score differences between predictions and targets:

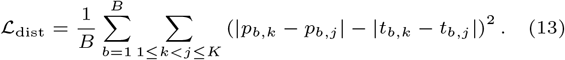

This term encourages the predicted score to respect not only the rank order but also the relative separation of decoys in quality space.

## Results and Discussion

### Evaluation metrics

We evaluate ABAG-rank against three baselines: (i) a random ranking baseline, (ii) the AF3 internal confidence-based ranking score, and (iii) DeepRank-Ab, a recently published graph neural network that leverages external HADDOCK-derived biophysical features. The performance of an antibody–antigen scoring model is evaluated based on two criteria: (i) its ability to rank the full ensemble in a way that is consistent with structural quality and (ii) its ability to retrieve the top*K* highest-quality decoys from an ensemble of predicted structures.

All results are reported on the hold-out test split, and for each complex, we restrict evaluation to a *strict intersection* of decoys for which *all* methods provide valid predictions. All metrics below are computed on this matched subset. This subset comprises 1091 complexes and 19,099 samples.

For a given complex, let *t*^*^ = max_*i*_ *t*_*i*_ denote the best achievable quality (DockQ) within the decoys pool. Each method assigns a score *ŝ*_*i*_ and thus induces a ranking. We denote by Top*K*(*ŝ*) the indices of the *K* highest-ranked decoys, and define the best recovered quality within the shortlist as

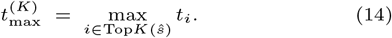

We report:

- **Tiered Top-***K* **Success (threshold-based)**. A method succeeds if the shortlist contains at least one decoy above a quality threshold *τ* :

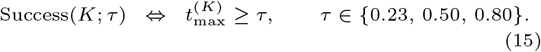

These thresholds correspond to acceptable, medium, and high-quality regimes of the DockQ metric as defined in [10].

- **Real Success (best-decoy retrieval)**. A stricter criterion that succeeds only if the method retrieves the *single* best decoy in the pool within the top-*K*:

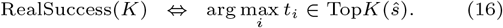
- **Relaxed Near-Optimal Success (gap-based)**. To capture the *usage perspective*, we also credit near-optimal retrieval when the selected decoy is equivalent to the best available structure. We quantify the quality gap as:

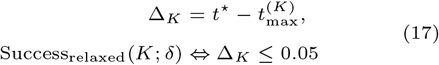

This metric distinguishes failure to retrieve the best model from cases in which the returned model lies within a small DockQ tolerance and is therefore equally valuable from a practical perspective. This metric is shown in Table 2 under the Real Success Rate (both for the best and the relaxed version).

#### Ranking correlation metrics

To evaluate the quality of the full ordering (beyond shortlist retrieval), we report:

- **Per-complex Spearman correlation**. For each complex *c*, we compute *ρ*_*c*_ = Spearman(*{ŝ*_*i*_*}, {t*_*i*_*}*), and summarize performance across complexes. This measures whether each method preserves relative order within an ensemble. While this is a standard metric, the per-complex Spearman correlation is especially relevant for complexes where the distribution of interface quality exhibits higher variance and is less relevant for complexes that cluster in a similar-quality region. (i.e., for a complex where all decoys fall in a cluster of very similar DockQ, the absolute correct ordering of the decoys is often irrelevant, while correct ordering for complexes where the variance of DockQ value of the decoys is much more important).
- **Global Spearman correlation**. We additionally compute the Global Spearman correlation after concatenating all matched decoys across complexes. This reflects overall monotonic agreement and is sensitive to cross-complex calibration.

The performance of ABAG-Rank model referred to throughout the main text includes the full set of input features as defined in the Methods.

### ABAG-Rank achieves the best performance at ranking interface quality

We first evaluate ABAG-Rank and the baseline methods on their ability to rank interface model quality at both global and per-complex levels. ABAG-Rank substantially outperforms both AF3 ranking confidence and DeepRank-Ab in global and per-complex Spearman correlation between predicted scores and ground-truth DockQ values while also predicting the interface quality with the lower numerical error (Table 1). Crucially, while being multiple orders of magnitude faster than DeepRank-Ab, ABAG-Rank shows a statistically significant improvement over AF3 confidence in per-complex Spearman correlation whereby DeepRank-Ab does not. AF3 and DeepRank-Ab tend to substantially overestimate the interface quality of models with incorrectly predicted binding pose, assigning high predicted scores when the ground-truth DockQ score is very low (Figure 3). In contrast, ABAG-Rank is much less prone to this failure mode and scores the interface quality more robustly, thereby reducing the number of high-scoring false positives. ABAG-Rank also achieves the highest AUC across classification thresholds defined by DockQ cutoffs (Acceptable > 0.23, Medium > 0.50, and High > 0.80), indicating stronger discrimination between low- and high-quality binding modes (Table 1 and Figure S6). When comparing the predicted ranking scores to the underlying ground-truth DockQ distribution, we observe that AF3 ranking confidence and DeepRank-Ab, indeed, fail to reproduce the shape of the true interface quality distribution (Figure S1). In contrast, ABAG-Rank most closely matches the ground-truth DockQ distribution among the evaluated methods. Notably, the internal AF3 confidence metrics pTM and ipTM exhibit a high density concentrated around very high score values, which might partly explain their tendency to assign overly confident scores even to low-quality models. Among the two, ipTM nevertheless more closely follows the overall shape of the ground-truth DockQ distribution.

**Table 1.** Performance of ABAG-Rank and baselines at ranking Ab-Ag models according to their interface quality. Best performance is indicated as bold. RMSE: Root Mean Sqaure Error, AUC: Area Under the Curve. Arrows down or up indicate the direction of preference. Statistical significance is denoted by ^ns^ (not significant) and ^***^ (*p <* 0.001). ^*†*^ for DeepRank-Ab denotes default inference settings of the published model. ^*^ for ABAG-Rank reports the time including the dataset preparation and the full inference pipeline with a batch size of 10.

| Method | Global Regression Metrics |  |  | AUC Classification (↑) |  |  | Inference |
| --- | --- | --- | --- | --- | --- | --- | --- |
|  | RMSE (↓) | Spearman (ρ) (↑) |  | Threshold (DockQ) |  |  | Time (s) (↓)<br>per Sample |
|  |  | Global | Per Cplx | Acc (> 0.23) | Med (> 0.50) | High (> 0.80) |  |
| Random | 0.453 | 0.007 | -0.010 | 0.500 | 0.502 | 0.510 | - |
| AlphaFold3 | 0.287 | 0.715 | 0.115 | 0.879 | 0.878 | 0.820 | - |
| DeepRank-Ab | 0.251 | 0.662 | 0.127<sup>ns</sup> | 0.857 | 0.872 | 0.850 | 23.805<sup>†</sup> |
| ABAG-Rank | **0.200** | **0.798** | **0.189<sup>\*\*\*</sup>** | **0.911** | **0.924** | **0.880** | **≤ 0.05<sup>\*</sup>** |

**Table 2.**
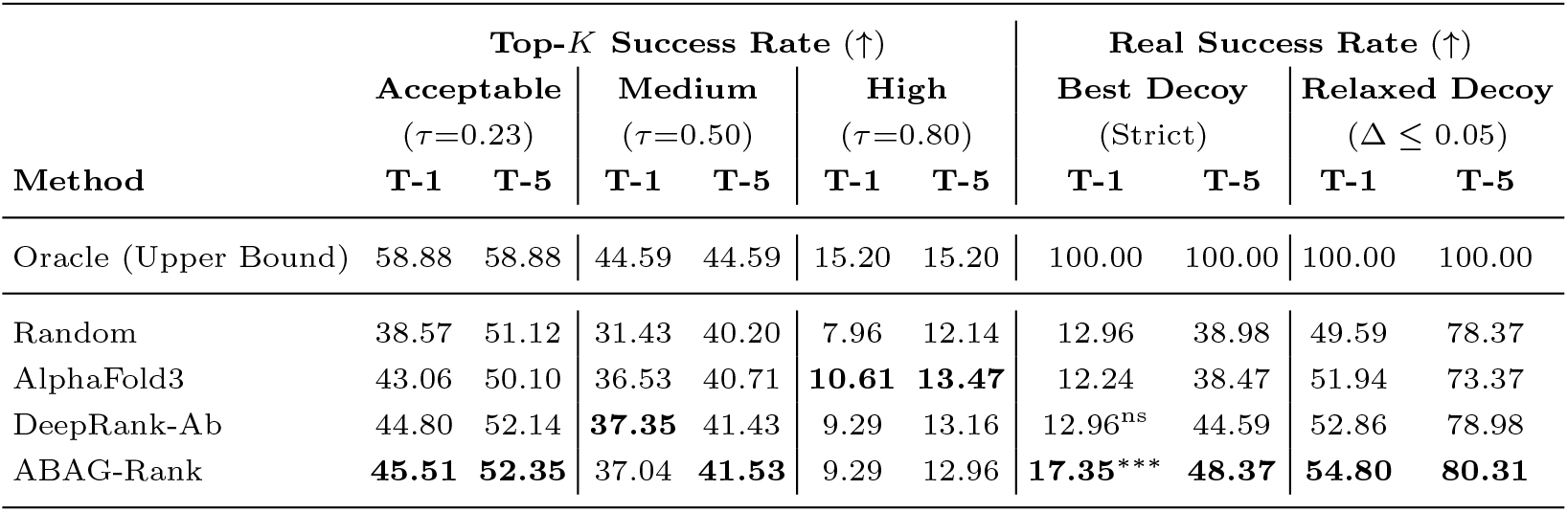
Performance of ABAG-Rank and baselines at selecting the best-interface quality Ab-Ag model among the top-*K* ranked samples. Best performance is indicated in bold. Statistical significance is denoted by ^ns^ (not significant), ^*^ (*p* < 0.05), ^**^ (*p <* 0.01), and ^***^ (*p* < 0.001).

| Method | Top-*K* Success Rate (↑) | | | | | | Real Success Rate (↑) | | | |
| --- | --- | --- | --- | --- | --- | --- | --- | --- | --- | --- |
| | Acceptable<br>($\tau=0.23$) | | Medium<br>($\tau=0.50$) | | High<br>($\tau=0.80$) | | Best Decoy<br>(Strict) | | Relaxed Decoy<br>($\Delta \leq 0.05$) | |
|  | T-1 | T-5 | T-1 | T-5 | T-1 | T-5 | T-1 | T-5 | T-1 | T-5 |
| Oracle (Upper Bound) | 58.88 | 58.88 | 44.59 | 44.59 | 15.20 | 15.20 | 100.00 | 100.00 | 100.00 | 100.00 |
| Random | 38.57 | 51.12 | 31.43 | 40.20 | 7.96 | 12.14 | 12.96 | 38.98 | 49.59 | 78.37 |
| AlphaFold3 | 43.06 | 50.10 | 36.53 | 40.71 | **10.61** | **13.47** | 12.24 | 38.47 | 51.94 | 73.37 |
| DeepRank-Ab | 44.80 | 52.14 | **37.35** | 41.43 | 9.29 | 13.16 | 12.96<sup>ns</sup> | 44.59 | 52.86 | 78.98 |
| ABAG-Rank | **45.51** | **52.35** | 37.04 | **41.53** | 9.29 | 12.96 | **17.35\*\*\*** | **48.37** | **54.80** | **80.31** |

**Figure 3.**
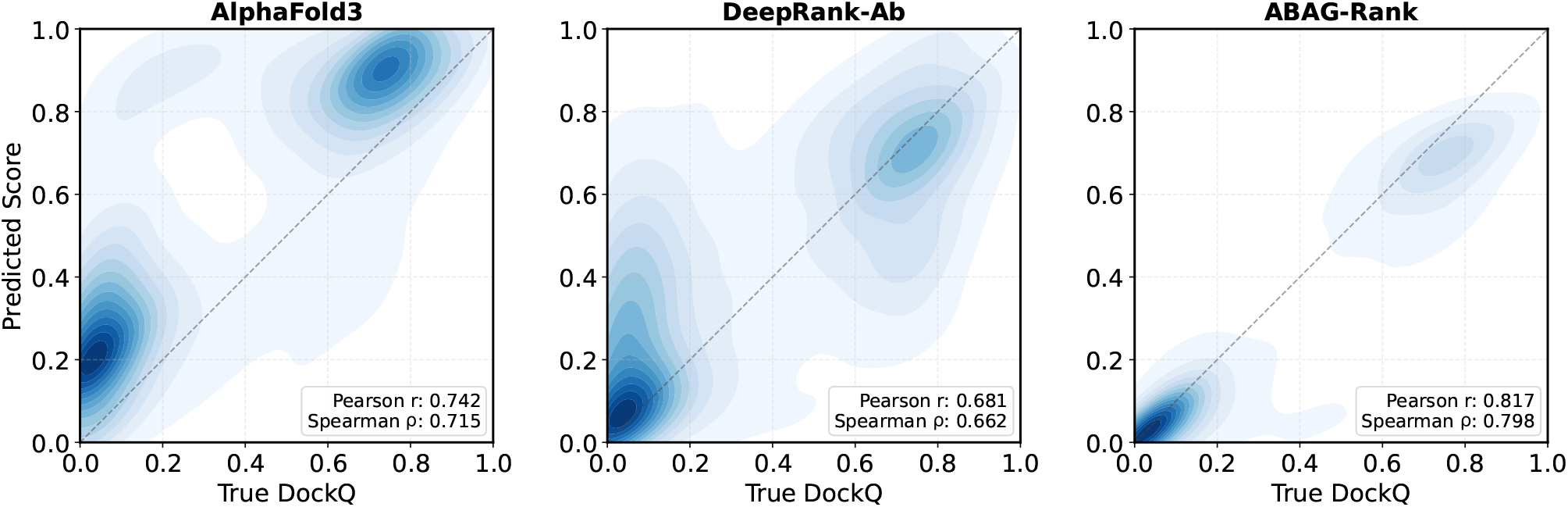
Distribution of predicted scores as a function of the ground-truth DockQ score for AlphaFold3 ranking confidence, DeepRank-Ab, and ABAGRank.Global Spearman and Pearson correlations are reported on the hold-out dataset comprised of 1091 Ab-Ag complexes not seen during training.

### ABAG-Rank improves retrieval of the best-available interface models

In practice, the most relevant objective is to retrieve the best available model of an Ab-Ag complex given a set of structural decoys, since ranking the predicted models alone does not necessarily lead to identification of the correct binding mode. We therefore evaluate whether ABAG-Rank can correctly identify the DockQ quality class of the top-*K* ranked samples and retrieve the sample with the highest possible interface quality within the top-*K* set. ABAG-Rank demonstrates either improved or on-par performance compared with AF3 ranking confidence and DeepRank-Ab in identifying *Acceptable* and *Medium* quality interfaces among the top-1 and top-5 ranked samples (Table 2). For the *High* quality class, however, AF3 is marginally better over the other methods.

When the task is defined more stringently as retrieving the best overall available interface-quality sample (Real Success), ABAG-Rank substantially outperforms both AF3 and DeepRank-Ab, although the success rate remains quite low: it correclty identifies the best sample in only 17.35% of cases at top-1 and 48.37% at top-5 level. Conceptually, retrieval of the *Best Decoy* is inherently challenging because it requires identifying the single numerically highest-DockQ sample, even when several alternative predictions could possess nearly identical interface and, consequently, numerically similar DockQ scores. Interestingly, when the criterion is therefore relaxed to allow a DockQ deviation of at most 0.05 from the best possible solution, the problem becomes substantially more tractable, indeed, reflecting the presence of many near-optimal samples clustered close to the top-scoring decoy. Under this relaxed definition, ABAG-Rank again achieves the best performance among all evaluated methods (Table 2).

This trend is further illustrated in Figure S2. In the top-*K* retrieval setting all methods progressively approach the Oracle upper bound as *K* increases, and by approximately top-10 they nearly recover a sample with the highest-quality interface, with ABAG-Rank performing marginally better than DeepRank-Ab and consistently better than AF3 ranking confidence. This indicates that, although the highest-quality solution is often not ranked first, it is typically found within a relatively small set of top-ranked candidates. In contrast, when considering only the single top-ranked prediction as a function of the sample pool size, all methods exhibit a clear performance saturation and remain consistently below the best available sample. This behavior is consistent with the previous observation that exact best-decoy retrieval is numerically stringent. Even as more samples become available, the ranking methods, including ABAG-Rank, struggle to reliably assign the optimal solution to the top-ranked decoy.

### Geometric features and AF3 confidence metrics are the key drivers of ABAG-Rank performance

To assess the contribution of individual input features to ABAG-Rank performance, we conducted an ablation study in which separate models were trained with specific input features removed. In particular, we evaluated the importance of ESM2 protein language model embeddings and the AF3-derived confidence metrics PAE and (i)pTM, which our full model relies upon.

For the overall best interface-quality model selection, we find that the model that uses only interface spatial *C*_*α*_ distances and one-hot residue identity encoding (i.e. without ESM, PAE and pTM), already improves over both AF3 ranking confidence and DeepRank-Ab by markedly reducing the number of high-scoring false positives (Figure S3). This suggests that simple structural and sequence-level descriptors might already be sufficient to provide a good initial estimate of Ab-Ag interface quality. Notably, DeepRank-Ab, despite relying on physics-derived features computed with HADDOCK, produces even a higher number of highly-scored incorrect predictions and underperforms the AF3 ranking confidence. The addition of (i)pTM and PAE to ABAG-Rank progressively improves its performance, while the inclusion of ESM embeddings yields the best overall performance (Figures S3, S6). Also, unlike AF3, ABAG-Rank does not show the tendency to assign similarly high scores to models with widely different interface qualities (Figure S5, complexes 8yor, 9dmq).

However, for the global model ranking, where the objective is to correctly order all samples by their interface quality, we observe that ABAG-Rank with ablated ESM, PAE, and ipTM features substantially underperforms the baselines. Markedly, the addition of ipTM alone is responsible for the largest improvement and brings the performance of ABAG-Rank above those of AF3 and DeepRank-Ab (Figure S4). We connect this observation with our earlier finding that ipTM resembles the ground-truth DockQ distribution (Figure S1) and therefore provides a reasonable initial estimate of interface model quality. Adding PAE further improves performance, and inclusion of ESM embeddings leads to the best overall result.

## Conclusions

This work introduces ABAG-Rank - a deep neural network for ranking interface quality of antibody-antigen models. ABAG-Rank achieves state-of-the-art performance on this task and outperforms existing baselines at both the global and per-complex levels, while also more effectively retrieving the best available structural solutions from AF3-generated ensembles. Importantly, the model addresses one of the key limitations of AF3 by substantially reducing the number of high-scoring false positives and assigning lower confidence to incorrect binding modes. Our results further show that simple geometric interface descriptors, when combined with AF3 internal confidence metrics and protein language model embeddings, provide a strong estimate of interface quality without requiring expensive physics-based calculations. In addition, ABAG-Rank is multiple orders of magnitude faster than another deep learning baseline DeepRank-Ab, making it well suited for large-scale evaluation of AF3 predictions in exploratory Ab-Ag docking studies. At the same time, the absolute performance of ABAG-Rank remains bounded by the generative capacity of AF3: if the sampled decoy pool does not contain high-quality conformations, no ranking model can recover the correct solution. Thus, while ABAG-Rank provides an effective solution to the ranking bottleneck, improving the generation of accurate candidate structures remains a fundamental challenge for the field.

## Author contributions statement

M.T. and M.L. prepared the dataset. M.T. developed the model with support and ideas from V.V., M.L., and A.E. V.V. and M.T. wrote the initial draft. All authors helped with analysis and writing. A.E. contributed funding.

## Acknowledgments

The authors thank anonymous reviewers for their valuable suggestions. AE was funded by Vetenskapsrådet, Grant No. 2021-03979, and the Knut and Alice Wallenberg Foundation, Grant No. 2022.0032. The computations and data handling were enabled by the supercomputing resource Berzelius, provided by the National Supercomputer Centre at Linköping University, the Knut and Alice Wallenberg Foundation, and SNIC, grant numbers SNIC 2021/5-297 and Berzelius-2021-29. The authors also express their gratitude to the rest of the Arne Elofsson lab for valuable discussions and input.

## Supplementary Information

### Supplementary Figures

**Fig. S1.**
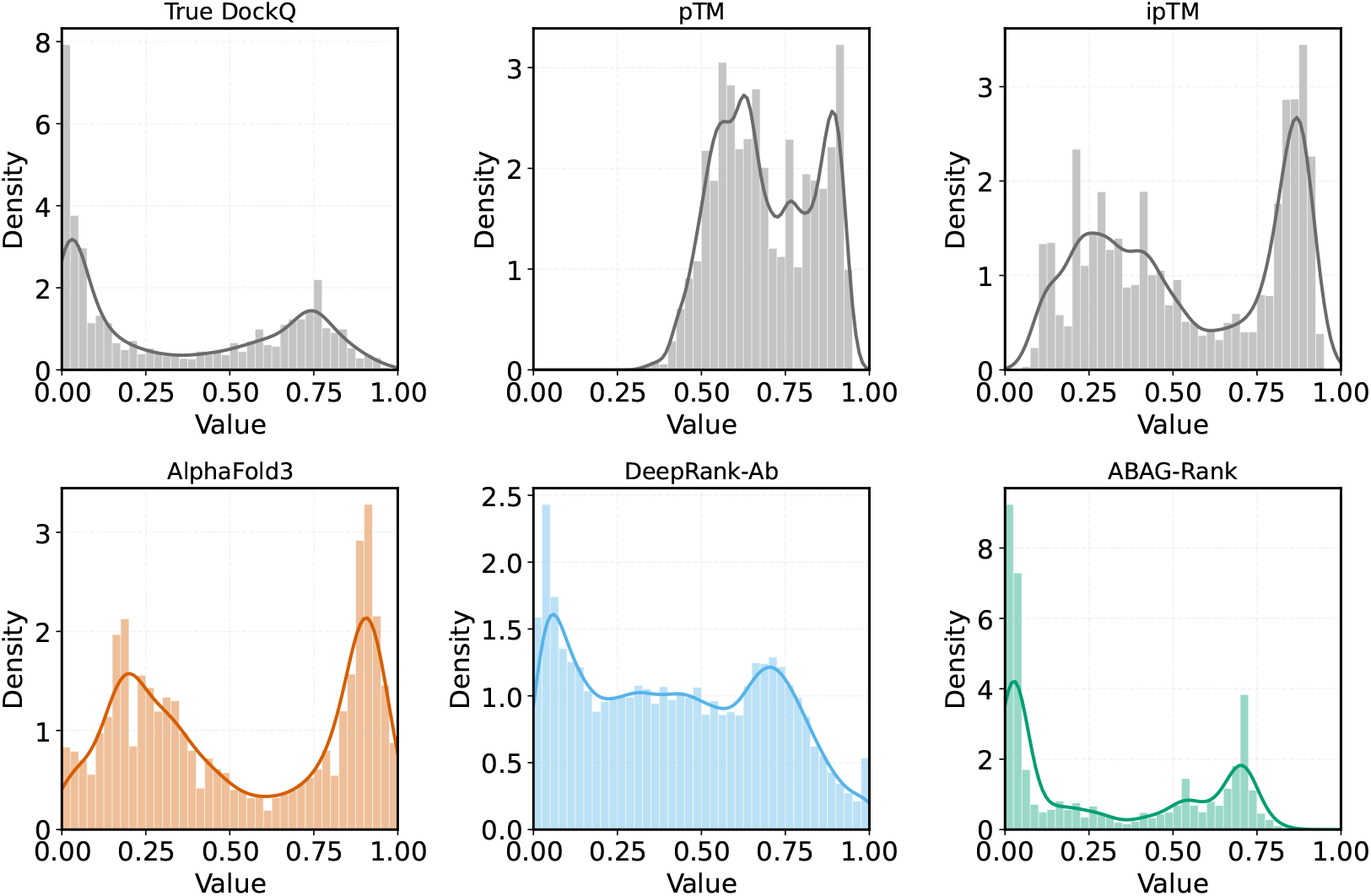
Distribution of the structural quality of AF3 predictions and the performance of the ranking methods along with pTM and iPTM for 1091 hold-out Ab-Ag complexes.

**Fig. S2.**
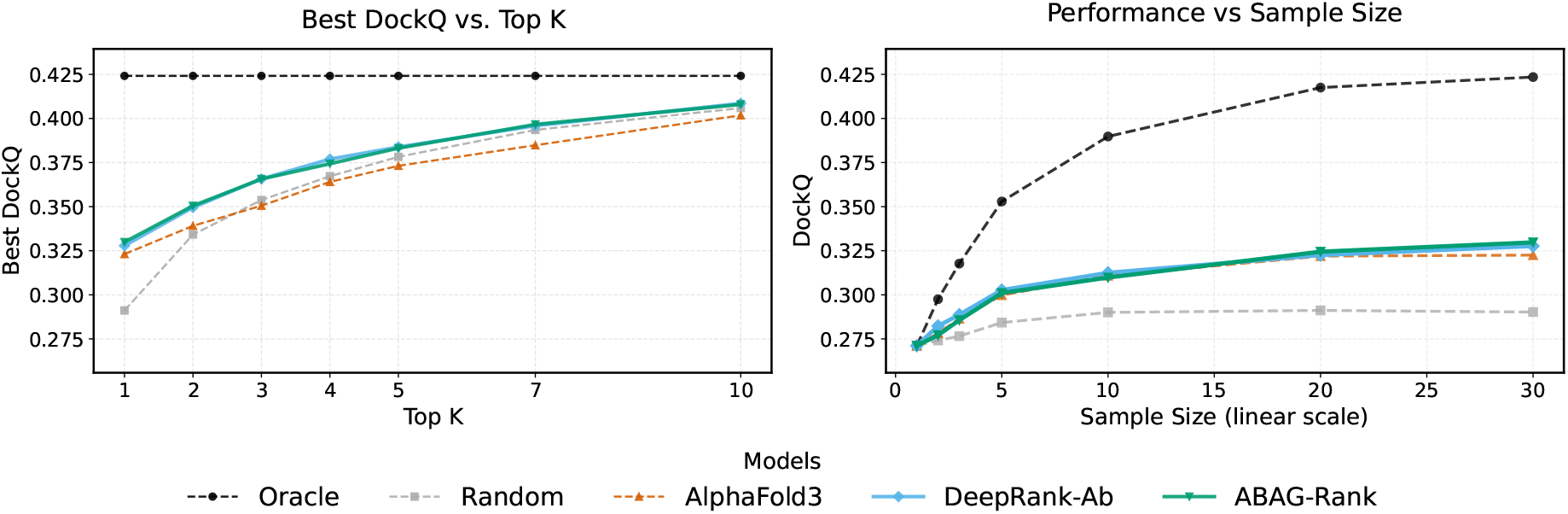
**Left:** Retrieval performance of the best-scoring model as a function of top-*K* models. DockQ is computed among the top-*K* models across the entire dataset comprised of 1091 hold-out complexes not seen during training. **Right:** Ranking performance as a function of decoy pool size, measured by the mean DockQ versus the sample set size *N* . The Oracle and Random curves denote the upper bound and the random-ranking baseline, respectively.

**Fig. S3.**
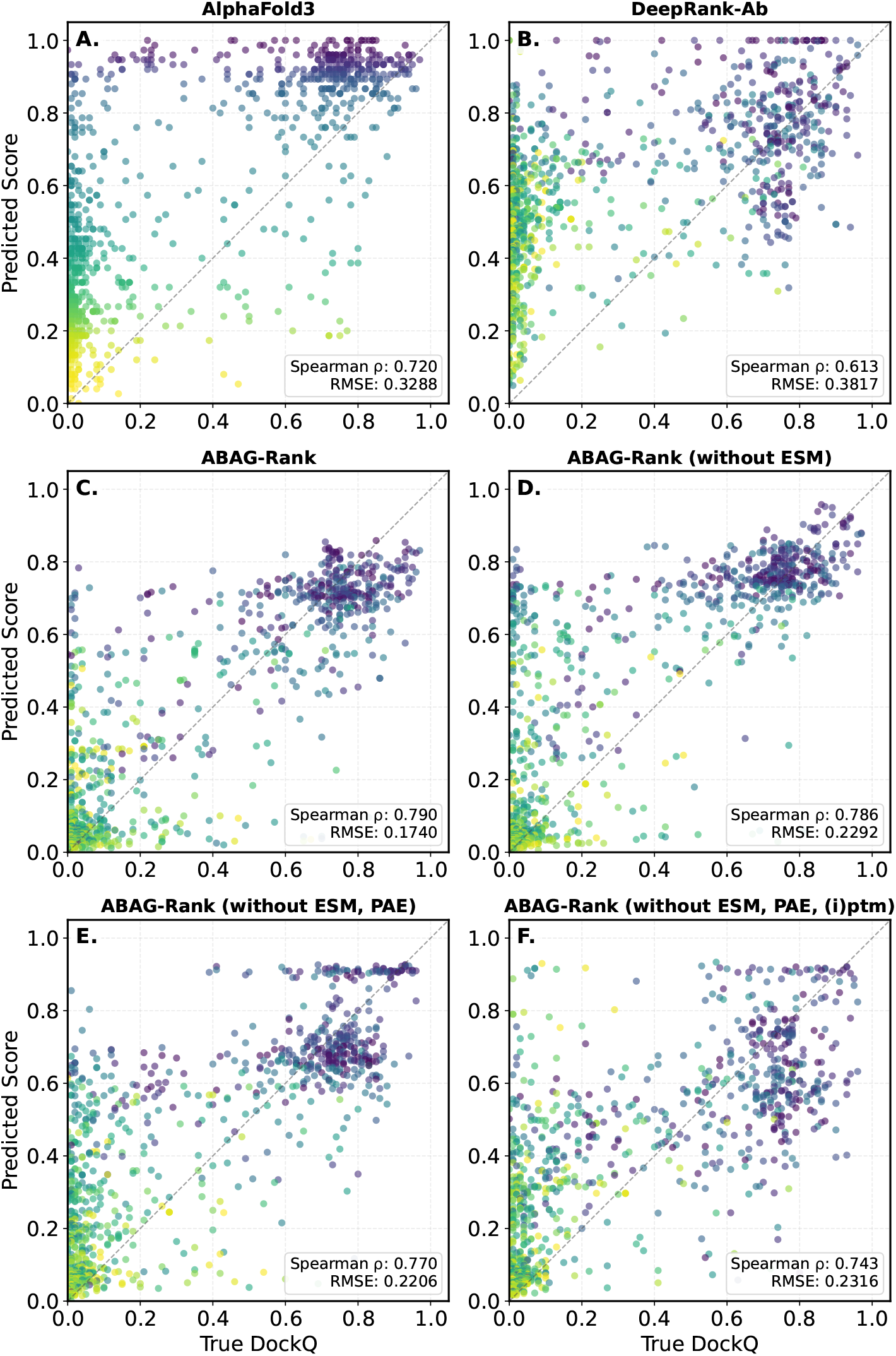
Comparison of ABAG-Rank and baseline models performance on the task of selecting the best-quality interface sample (top-1) for 1091 hold-out complexes. (C) Denotes the models reported in the main text which relies upon ESM, PAE and (i)pTM features.

**Fig. S4.**
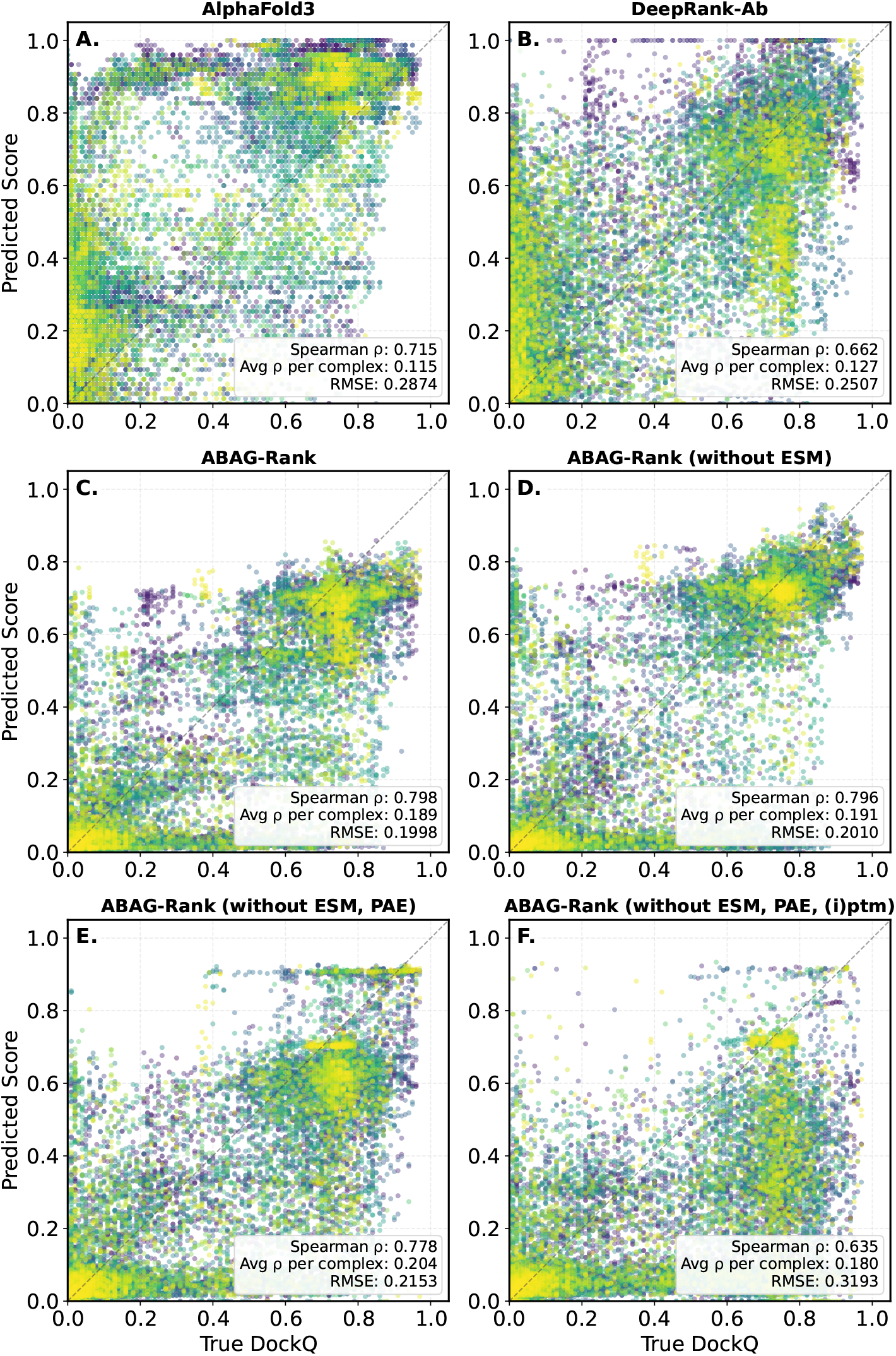
Comparison of ABAG-Rank and baseline models on global ranking of Ab-Ag models according to their interface quality for all 19,099 samples across 1091 hold-out complexes. (C) Denotes the models reported in the main text which relies upon ESM, PAE and (i)pTM features.

**Fig. S5.**
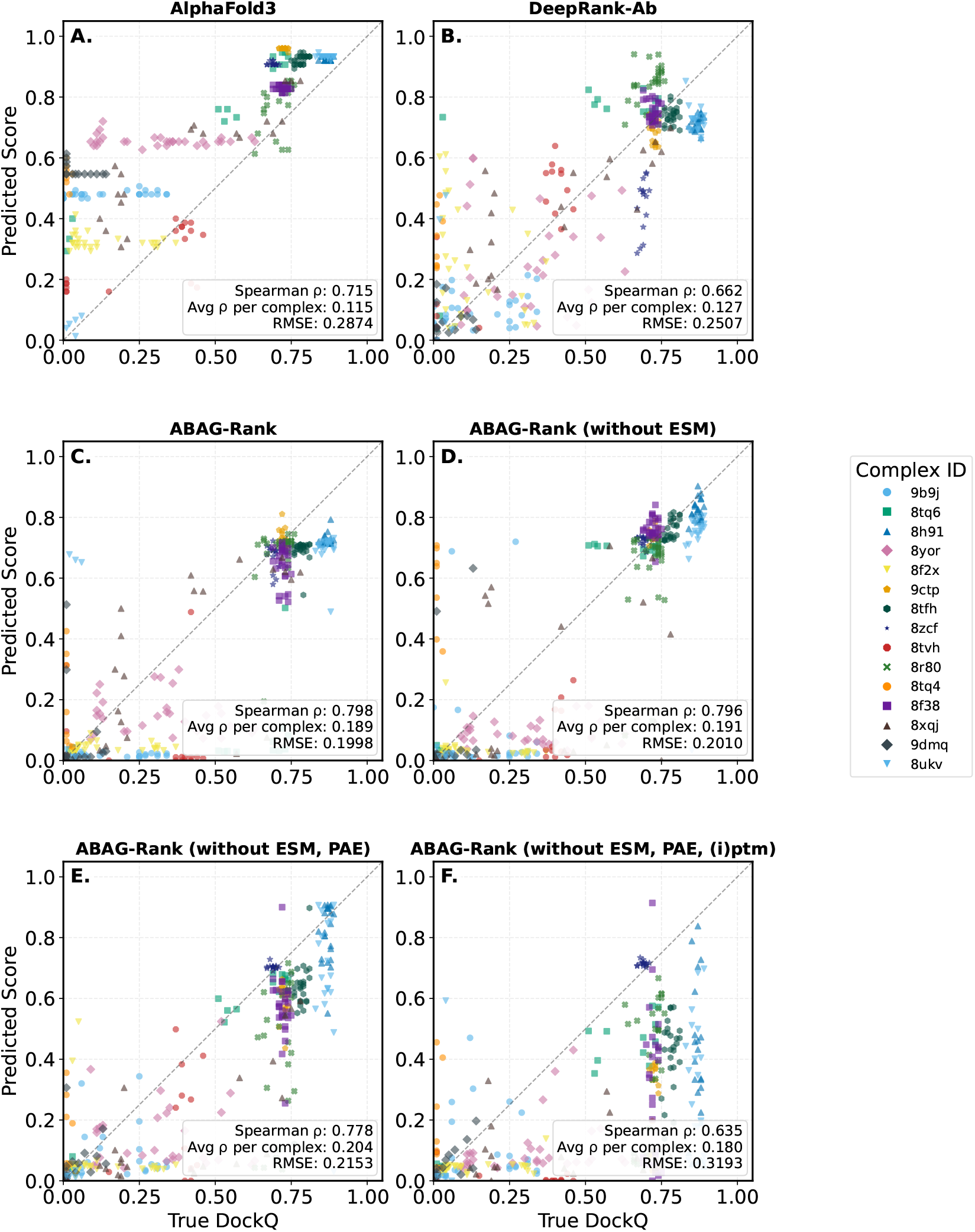
Complex-Specific Performance Examples. Scatter plots of a random subset of Complex ID and the corresponding within-complex ranking.

**Fig. S6.**
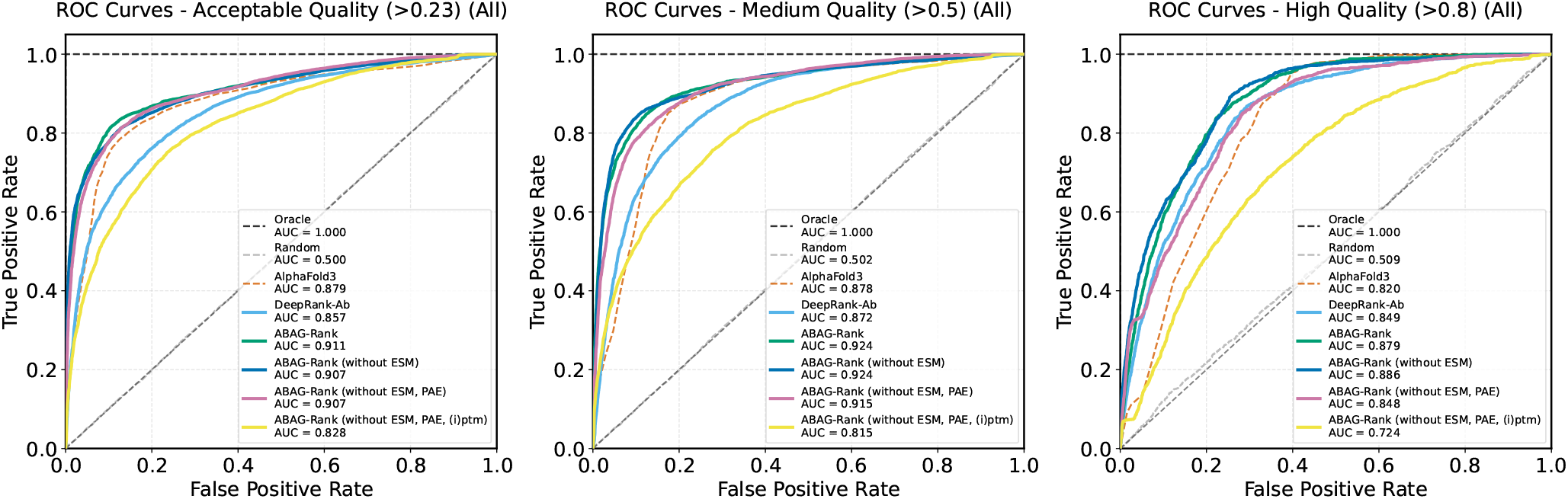
ROC Curves of ABAG-Rank and baseline models across DockQ quality thresholds computed on the hold-out set of 1091 complexes not seen during training. Ablated features for ABAG-Rank are indicated in the figure legend.

### Dataset Filtering Algorithms

#### Algorithm 1

IsRedundant

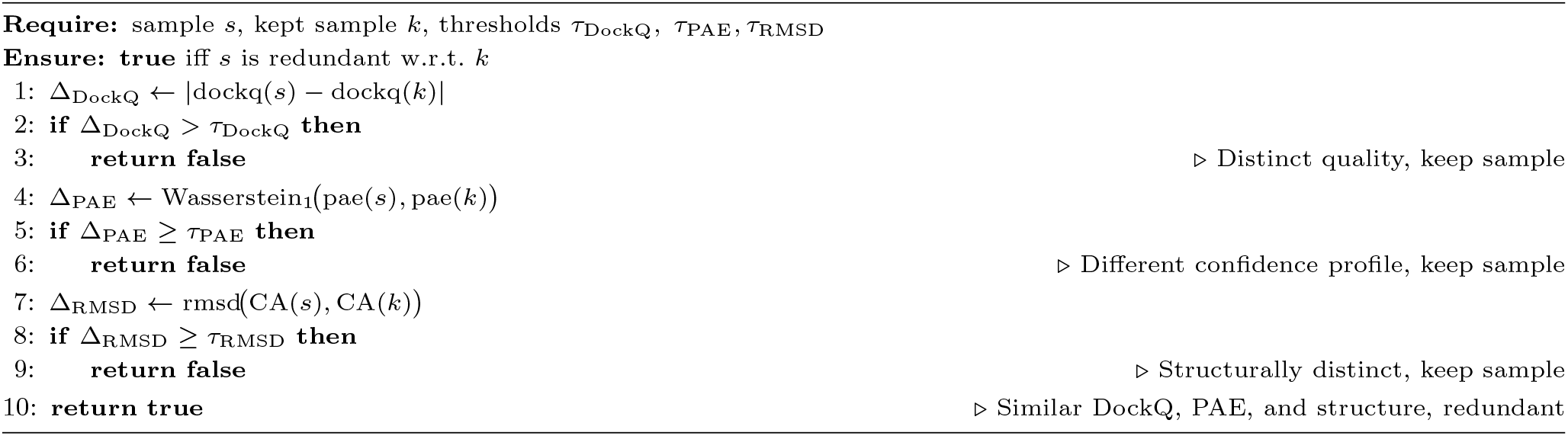

#### Algorithm 2

GreedyPrune

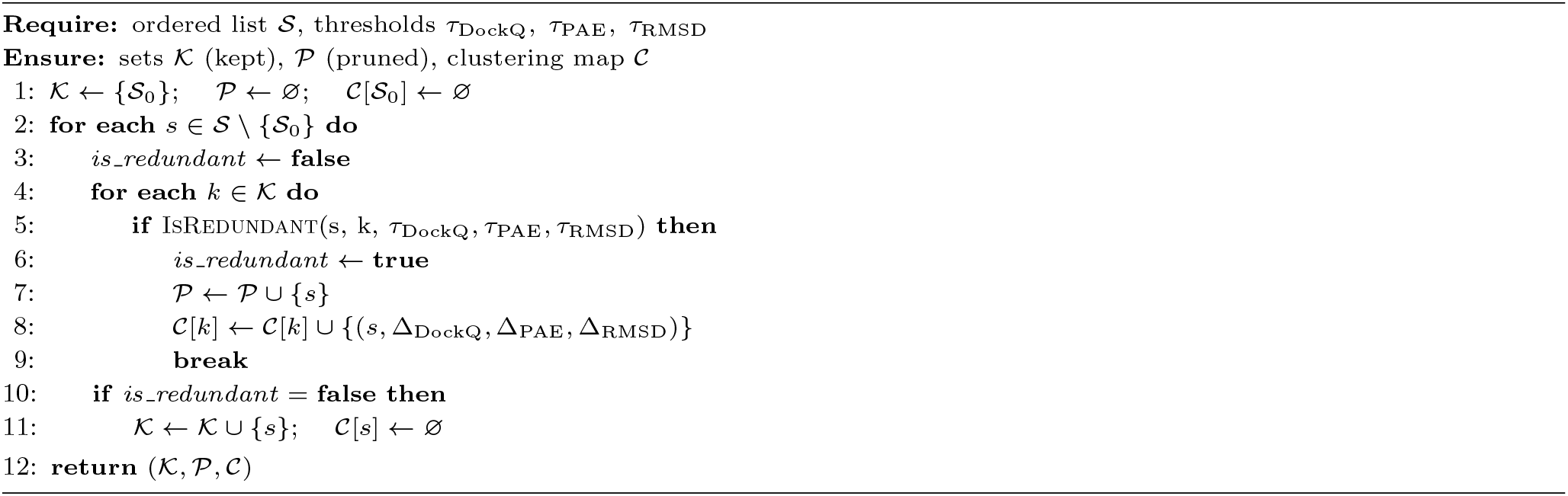

## References

1. John Jumper, Richard Evans, Alexander Pritzel, Tim Green, Michael Figurnov, Olaf Ronneberger, Kathryn Tunyasuvunakool, Russ Bates, Augustin Žídek, Anna Potapenko, Alex Bridgland, Clemens Meyer, Simon A. A. Kohl, Andrew J. Ballard, Andrew Cowie, Bernardino Romera-Paredes, Stanislav Nikolov, Rishub Jain, Jonas Adler, Trevor Back, Stig Petersen, David Reiman, Ellen Clancy, Michal Zielinski, Martin Steinegger, Michalina Pacholska, Tamas Berghammer, Sebastian Bodenstein, David Silver, Oriol Vinyals, Andrew W. Senior, Koray Kavukcuoglu, Pushmeet Kohli, and Demis Hassabis. Highly accurate protein structure prediction with AlphaFold. Nature, 596(7873):583–589, August 2021.

2. Josh Abramson, Jonas Adler, Jack Dunger, Richard Evans, Tim Green, Alexander Pritzel, Olaf Ronneberger, Lindsay Willmore, Andrew J. Ballard, Joshua Bambrick, Sebastian W. Bodenstein, David A. Evans, ChiaChun Hung, Michael O’Neill, David Reiman, Kathryn Tunyasuvunakool, Zachary Wu, Akvilė Žemgulytė, Eirini Arvaniti, Charles Beattie, Ottavia Bertolli, Alex Bridgland, Alexey Cherepanov, Miles Congreve, Alexander I. Cowen-Rivers, Andrew Cowie, Michael Figurnov, Fabian B. Fuchs, Hannah Gladman, Rishub Jain, Yousuf A. Khan, Caroline M. R. Low, Kuba Perlin, Anna Potapenko, Pascal Savy, Sukhdeep Singh, Adrian Stecula, Ashok Thillaisundaram, Catherine Tong, Sergei Yakneen, Ellen D. Zhong, Michal Zielinski, Augustin Žídek, Victor Bapst, Pushmeet Kohli, Max Jaderberg, Demis Hassabis, and John M. Jumper. Accurate structure prediction of biomolecular interactions with AlphaFold 3. Nature, 630(8016):493–500, June 2024.

3. Samuel Fromm, Marko Ludaic, and Arne Elofsson. Evaluating deep learning based structure prediction methods on antibody-antigen complexes. bioRxiv, 2025.

4. Eva Smorodina, Montader Ali, Klara Kropivsek, Leonardo Salicari, Samo Miklavc, Aibek Kappassov, Chengcheng Fu, Pietro Sormanni, Ario de Marco, and Victor Greiff. Structural plausibility without binding specificity: Limits of ai-based antibody-antigen structure prediction confidence scores. bioRxiv, pages 2026–03, 2026.

5. Mu Gao and Jeffrey Skolnick. Improved deep learning prediction of antigen-antibody interactions. Proceedings of the National Academy of Sciences, 121(41):e2410529121, 2024.

6. Björn Wallner. Afsample: improving multimer prediction with alphafold using massive sampling. Bioinformatics, 39(9):btad573. 09 2023.

7. Nessim Raouraoua, Claudio Mirabello, Thibaut Véry, Christophe Blanchet, Björn Wallner, Marc F. Lensink, and Guillaume Brysbaert. MassiveFold: unveiling AlphaFold’s hidden potential with optimized and parallelized massive sampling. Nature Computational Science, 4(11):824–828, November 2024.

8. Patrick Bryant, Gabriele Pozzati, and Arne Elofsson. Improved prediction of protein-protein interactions using alphafold2. Nature communications, 13(1):1265, 2022.

9. Wensi Zhu, Aditi Shenoy, Petras Kundrotas, and Arne Elofsson. Evaluation of alphafold-multimer prediction on multi-chain protein complexes. Bioinformatics, 39(7):btad424, 2023.

10. Sankar Basu and Björn Wallner. Dockq: A quality measure for protein-protein docking models. PLOS ONE, 11(8):1–9, 08 2016.

11. Julia K Varga, Sergey Ovchinnikov, and Ora Schueler-Furman. actifptm: a refined confidence metric of alphafold2 predictions involving flexible regions. Bioinformatics, 41(3):btaf107. 03 2025.

12. Roland L Dunbrack Jr. Rēs ipsae loquunt: What’s wrong with alphafold’s iptm score and how to fix it. bioRxiv, 2025.

13. Matthew McFee, Jisun Kim, and Philip M Kim. Eudockscore: Euclidean graph neural networks for scoring protein–protein interfaces. Bioinformatics, 40(11):btae636. 10 2024.

14. Matthew McFee and Philip M Kim. Gdockscore: a graph-based protein–protein docking scoring function. Bioinformatics advances, 3(1):vbad072, 2023.

15. Xiaotong Xu, Ilaria Coratella, Victor Reys, and Alexandre MJJ Bonvin. Deeprank-ab: a scoring function for antibody-antigen complexes based on geometric deep learning. bioRxiv, pages 2025–12, 2025.

16. Marco Giulini, Victor Reys, João MC Teixeira, Brian Jiménez-García, Rodrigo V. Honorato, Anna Kravchenko, Xiaotong Xu, Raphaëlle Versini, Anna Engel, Stefan Verhoeven, et al. Haddock3: a modular and versatile platform for integrative modeling of biomolecular complexes. Journal of Chemical Information and Modeling, 65(13):7315–7324, 2025.

17. Manzil Zaheer, Satwik Kottur, Siamak Ravanbakhsh, Barnabas Poczos, Russ R Salakhutdinov, and Alexander J Smola. Deep sets. In I. Guyon, U. Von Luxburg, S. Bengio, H. Wallach, R. Fergus, S. Vishwanathan, and R. Garnett, editors, Advances in Neural Information Processing Systems, volume 30. Curran Associates, Inc., 2017.

18. Zeming Lin, Halil Akin, Roshan Rao, Brian Hie, Zhongkai Zhu, Wenting Lu, Nikita Smetanin, Robert Verkuil, Ori Kabeli, Yaniv Shmueli, Allan dos Santos Costa, Maryam Fazel-Zarandi, Tom Sercu, Salvatore Candido, and Alexander Rives. Evolutionary-scale prediction of atomic-level protein structure with a language model. Science, 379(6637):1123–1130, 2023.

19. James Dunbar, Konrad Krawczyk, Jinwoo Leem, Terry Baker, Angelika Fuchs, Guy Georges, Jiye Shi, and Charlotte M. Deane. SAbDab: the structural antibody database. Nucleic Acids Research, 42(D1):D1140–D1146, November 2013.

